# Dynamic characteristics and Functional Analysis Provide new insights into the role of *GauERF105* in resistance against Verticillium wilt in Cotton

**DOI:** 10.1101/2021.12.30.474616

**Authors:** Yanqing Wang, Muhammad Jawad Umer, Yuqing Hou, Yanchao Xu, Teame Gereziher Mehari, Jie Zheng, Yuhong Wang, CAI, Zhongli Zhou, Zhikun Li, Fang Liu

## Abstract

Verticillium wilt is the most devastating disease of cotton and it results in huge yield losses every year in the fields. The underlying mechanisms of VW in cotton are not well explored yet. In the current approach we used the transcriptome data from *G. australe* in response to Verticillium wilt attack to mine the ERF TFs and prove their potential role in resistance against VW attack in cotton. We identified 23 ERFs in total, and on the basis of expression at different time points i.e., 24h, 48h and 72h post inoculation and selected *GauERF105* for further validation. We performed VIGS in cotton and over expression in Arabidopsis respectively. Moreover, DAB and trypan staining also suggests that the impact of disease was more in the wildtype as compared to transgene lines. On the basis of our results, we confirmed that *GauERF105* is the key candidate and playing a key role for defending cotton against VW attack. Current finding might be helpful for generating resistance germplasm in cotton and it will be beneficial to recover the yield losses in field.

## 1. Introduction

Cotton is an economic crop, while verticillium wilt severely restricts cotton production (T. Li, Zhang, Jiang, Li, & Dhar, 2021). Cotton is one of the species of Malvaceae with high economic value and wide geographical distribution. There are a total of 53 species of cotton (Kunbo, Wendel, & Jinping, 2018), of which 46 diploid cotton species divided into 8 genome groups of A, B, C, D, E, F, G, and K. The remaining 7 tetraploid cotton species (AD)_1_-(AD)_7_ belong to one allopolyploid genome(Beasley, 1942; Fryxell, 1992; Phillips, 1966). The A, B, E and F cotton type genomes are distributed in Asia and Africa, the C, G and K cotton type genomes are distributed in Australia, and the D and AD cotton type genomes are distributed in the America.

Verticillium wilt can infect the vascular bundles of cotton plant. Verticillium wilt was first discovered in the United States in 1915, and then introduced to China in 1935, and successively broke out in major cotton production areas (Jian, Lu, Xiu, Wang, & Zhang, 2004). *Verticillium dahliae* is the pathogen of cotton verticillium wilt, which is mainly divided into *Verticillium dahlia* and *Verticillium albo-atrum. Verticillium dahliae* can cause the wilt of a variety of plants. In addition to cotton, fruits, and vegetables such as potatoes, tomatoes, grapes, and some woody plants can be attacked by *Verticillium dahliae* (Barbara, 2003). The main host of Verticillium black and white are alfalfa, hops, soybeans, tomatoes, potatoes, and some weeds (Chen, Lee, & Robb, 2004; Ligoxigakis, Vakalounakis, & Thanassoulopoulos, 2002). After investigations, the diseases in cotton fields in China are mainly caused by *Verticillium dahliae* (Yinhua et al., 2014).

Cotton verticillium wilt is one of the soil-borne fungal vascular disease, which leads to yearly yield losses of over 30% and a severe economic loss of roughly 250– 310 million dollars for China (Gong et al., 2017). Though, it is hard to control pathogenic harm to cotton plants, despite the fact that many attempts were made, among them are the use of fungicides as well as cultural methods (Mohamed & Akladious, 2017; Wei & Yu, 2018). Generally, protecting plants from pathogenic damage, disease-resistant cultivars must be widely planted. Although certain genes/proteins/TFs were characterized in cotton plant protection against pathogens, several effective candidate genes were updated for their use in disease-resistant breeding (Cai et al., 2009; Gong et al., 2017; Zeng, Chen, Luo, & Tian, 2016). Thus, molecular mechanisms of plant resistance against the *V. dahliae* as well as the functional analysis of genes linked to defense must be explored deeply.

The ERF transcription factor contains an AP2 domain, which is composed of a transcription regulatory domain, a DNA domain, and a nuclear localization sequence (NLS). In addition, some ERF family members contain oligomerization sites and phosphoric acid modification sites to regulate gene expression. At the N-terminus of the ERF domain is a basic hydrophilic region, which contains 3 β-sheet structures, in which the 14^th^ alanine and 19^th^ aspartic acid residues in the second β-sheet helps in binding ERF transcription factors to different cis-acting elements (Stockinger, Gilmour, & Thomashow, 1997). DRE / CRT (Dehydration response element, DRE; C-repeat, CRT) and GGC-box are the main cis-acting elements that ERF binds with. Among them, DRE/CRT is mainly related to abiotic stress, and GCC-box is mainly involved in regulating biotic stress. The promoter region of many drought response-related genes contains a large amount of DRE, and its core sequence is TACCGACAT (Kizis & Pagès, 2002). The core sequence TGGCCGAC of CRT element is mainly present in low temperature response genes. It is precisely because the cis-elements of DRE/CRT have the core sequence CCGAC, which is usually related to temperature, salt, and drought, so they are usually referred to as the cis-acting elements of DRE/CRT (Fujimoto, Ohta, Usui, Shinshi, & Ohme-Takagi, 2000; Kizis & Pagès, 2002). GCC-box has a conservative AGCCGCC sequence, which is generally present in the promoter regions of many disease-related protein genes. ERF can respond to some biological stress responses by combining with GCC-box (Meng et al., 2010; H.-J. Yang et al., 2002).

Among the signal pathways that plants respond to biological stress, the ethylene signaling pathway plays an important role, and many disease-resistant genes are induced and regulated by this signal pathway (C. Yang, Lu, Ma, Chen, & Zhang, 2015). Overexpression of *AtERF1* directly activates the expression of plant defensins (PDF1.2) and improves plant resistance to pathogens (Berrocal - Lobo, Molina, & Solano, 2002; Lorenzo, Piqueras, S á nchez-Serrano, & Solano, 2003). The T-DNA insertion mutant of *AtERF14* increases the susceptibility of Arabidopsis to *Fusarium oxysporum* infection (Oñate-Sánchez, Anderson, Young, & Singh, 2007). *AtERF96* positively regulates the resistance of Arabidopsis to necrotic pathogens by enhancing the expression of PDF1.2a, PR-3, PR-4, and ORA59 (Catinot et al., 2015). The overexpression of *GmERF5* in soybean increases its resistance to *Phytophthora sojae*, and it can positively regulate the expression of PR genes after being induced by *Phytophthora sojae* (L. Dong et al., 2015). Zang et al. found that overexpression of *ZmERF105* can increase the resistance of maize to *S. sphaerocephala*, while the *erf105* mutant strain showed the opposite phenotype; after *ZmERF105* overexpression strains were infected with *S. sphalacca, ZmPR1a, ZmPR2, ZmPR5, ZmPR10*.*1*. The expression of disease-related genes such as *ZmPR10*.*2* is enhanced, on the contrary, the expression of PR gene is reduced in the ERF105 mutant line (Zang et al., 2020). Meng et al. cloned two ERF transcription factor members *EREB1* and *EREB2* from sea island cotton, and *Verticillium dahliae* can induce the expression of these two genes (Meng et al., 2010). Guo et al. used the SSH method to enrich some differentially expressed genes related to defense response and cloned the gene *GbERF1-like* from sea island cotton. The study showed that the overexpression of *GbERF1-like* activated the synthesis of lignin-related genes, and enhanced cotton and pseudo-resistance of Arabidopsis to Verticillium wilt (Guo et al., 2016).

Here we, cloned *GauERF105* gene from the *G. australe* and then verified the gene function in cotton via VIGS and overexpression in Arabidopsis in response to VW attack. This research will provide a basis for further mining of excellent disease resistance genes in wild cotton, in-depth research on the molecular mechanism of cotton resistance to Verticillium wilt and provide new genetic resources for cotton disease resistance breeding.

## 2 Materials and methods

### 2.1 Plant material, *Verticillium dahliae* strains and gene selection

The Zhongzhimian 2 (disease-resistant upland cotton variety), Diploid wild cotton (*G. australe*) and the Colombian ecotype Arabidopsis were provided by the Cotton Research Institute of the Chinese Academy of Agricultural Sciences, Anyang, China. The cotton seedlings are grown in a growth box with a light/dark cycle of 16/8h and a temperature of 27°C (day)/23°C (night). The wild-type *Arabidopsis thaliana* (COL-0) was cultured under a light/dark cycle of 16/8h at a constant temperature of 22°C. The highly invasive strain of *V. dahliae* (LX2-1) was used for disease resistance identification, and the preparation of conidia suspension (10^7^conidia mL^-1^ for cotton, 10^6^ and 10^3^ conidia for Arabidopsis) and inoculation are as described above [1]. From previously available RNA-Seq data(Q. Dong et al., 2019), we searched all the ERF family genes and on the basis of expression we selected *GauERF105* for further experiments.

### 2.2 Gene cloning and Phylogenetic Analysis

RNAprep Pure Plant Plus Kit (TIANGEN BIOTECH, Beijing, China) was used to extract the sample RNA, and the quality of the sample was checked by agarose gel electrophoresis and spectrophotometer. TranScript-All-in-One First-Strand cDNA Synthesis SuperMix (TransGen, Beijing, China) reverse transcription kit was used to obtain the cDNA. Design primers based on the CDS sequence of *GauERF105*, use G. australe cDNA as a template, and use P505 high-fidelity polymerase (Vazyme, Nanjing, China) to amplify the target gene. Download the amino acid sequences of other cotton ERF members from the NCBI website. DNAMAN software was used for multiple sequence alignment, and MEGA-X was used to construct a phylogenetic tree.

### 2.3 Cotton VIGS and quantification of disease resistance

Virus induced gene silencing (VIGS) was performed according to the procedure described previously by (Q. Dong et al., 2019). A 432 bp *GhERF105* fragment was amplified and inserted between the BamHI and EcoRI sites of the tobacco Rattle virus (TRV) binary vector pTRV2. Phytoene desaturase (PDS) gene was used as a marker to detect the reliability of silencing. These experiments were repeated three times independently, using more than 35 plants for each treatment. At 25dpi, the seedlings symptoms are divided into five levels: 0, 1, 2, 3, and 4 according to the symptoms on the leaves (Z. K. Li et al., 2019). The calculation of Disease index (DI) was as follows: **DI = [(Σdisease grades × number of infected plants) / (total number of scored plants × 4)] ×100 (Cai et al**., **2020)**.

### 2.4 Generation of transgenic Arabidopsis lines

We used the method of homologous recombination to link the gene with ‘BamHI’ and ‘SacI’ restriction sites with the overexpression vector PBI121 to obtain the expression vectorPBI121-GauERF105, which was then transformed into *Agrobacterium tumefaciens* GV3101. Transgenic *Arabidopsis thaliana* plants were obtained using the flower soaking method (Clough & Bent, 1998). The transgenic lines (T0, T1 and T2 seeds) were screened on half-strength MS medium with kanamycin added. The T3 transgenic lines were identified and characterized by qRT-PCR and then used in subsequent experiments. The 20-day-old Arabidopsis plants were inoculated with *V. dahliae*. Twenty days after inoculation (20Dpi), the symptoms were scored. According to the degree of leaf yellowing, the degree of resistance to VW is graded from 0 to 4. The calculation method of the disease index was kept same as above.

### 2.5 Histochemical staining of cotton stem lignin

The Wiesner method (Speer, 1987) was used to analyze the lignin histochemical staining of cotton. The parts of cotton cotyledon nodes of wild-type, TRV: 00 and TRV: *GhERF105* plants were sectioned by hand. Dip the slices with Wiesner reagent [3% (w/v) phloroglucinol in dd solution, solubilized with absolute ethanol] for 5 minutes, wash twice with distilled water, acidify with 6% hydrochloric acid solution for 5 minutes, and wash away residual after hydrochloric acid treatment, place it on a glass slide to observe and take pictures under a stereo microscope.

### 2.6 Fungal recovery assay of cotton stems after *V. dahliae* inoculation

We performed the fungal recovery assay as described earlier by (Song & Thomma, 2018).We randomly took cotton plants treated with Verticillium wilt. We used the stem sections that were above the cotyledons and placed them in a sterilized triangular flask and use disinfectant to disinfect the surface of the cotton stems for 7 minutes and sterilize them immediately after disinfection. Wash the stem with ddH_2_O, rinse 3 times for 5min each time. Place the samples in a petri dish containing PDA with cephalosporin and incubate at 25°C in the dark for 3-5 days then observe the fungal growth.

### 2.7 DAB staining

The 3,3’diaminobiphenyl (DAB) staining method as described by (Gao et al., 2013) for estimating the production and accumulation of hydrogen peroxide in the leaves. After 72 hours of inoculation with *Verticillium dahliae*, Arabidopsis leaves were taken, rinsed with distilled water, and then dried with filter paper. Add the leaves in a 2mL centrifuge tube, take an appropriate amount of DAB staining solution for staining, and store in the dark at room temperature for 8h. Remove the staining solution, add 95% ethanol to remove chlorophyll, keep changing ethanol for 2-3 days. Use sterilize water to wash the leaves before taking pictures.

### 2.8 Trypan blue staining

The true leaves of the *GauERF105* Arabidopsis experimental group and the wild-type Arabidopsis WT blank control group were respectively inoculated for 72 hours and soaked in trypan blue dye solution (10 mL lactic acid, 10 mL glycerin, 10 g phenol, 10 mg Trypan blue, 10 mL of distilled water), in a boiling water bath for 2 minutes, decolorize in chloral hydrate (2.5 g/mL) after cooling, replace the decolorizing solution every day, decolorize for 3 days, then wash with sterile water, take pictures and record.

### 2.9 Expression analysis of defense marker genes

Using cotton and *Arabidopsis thaliana* inoculated with *Verticillium dahliae* as materials, leaf tissues were obtained at 48hpi and 72hpi respectively, and then quickly frozen in liquid nitrogen to extract total RNA. These defensive marker genes in Arabidopsis and cotton were detected using specific primers of some disease-related proteins (PRs) described by (Guo et al., 2016). Each sample has 3 biological replicates and 3 technical replicates.

### 2.10 qRT-PCR analysis

According to the above method, the total RNA of the plant was extracted and reverse transcribed into cDNA. PCR amplification was used SYBR®qPCR Master Mix (Vazyme, Nanjing, China) The cotton GhUBQ7 gene and the Arabidopsis AtACTIN gene were used as internal reference genes for qRT-PCR analysis, Calculate the relative expression of genes according to the 2^-ΔΔct^ method. The qRT-PCR assay were performed as described previously

## 3 Results

### 3.1 Cloning and Sequence Analysis of *GauERF105*

Based on already published RNA-seq data(Q. Dong et al., 2019) of *G. australe* under *V. dahliae* treatment, we identified all the ERF family genes. On the basis of gene expression analysis, we selected a gene namely *GauERF105* for subsequent analysis (Figure 1A). The CDS sequence of *GauERF105* consisting of 639bp was successfully cloned from the *G. australe*. This gene encodes 213 amino acids with theoretical molecular mass of 23.609kD and an isoelectric point of 8.99. *GauERF105* was found to be located on chromosome 12 (10153228-10153866). Blastp search on NCBI using the protein sequences of *GauERF105* revealed that it has a similarity to *Gossypium raimondii* (XP_01243908.1), *Gossypium arboretum* (XP_017636134.1), *Gossypium hirsutum* (XP_016721164.1), *Theobroma cacao* (XP_017873866.1), *Herrania umbratical*(XP_021286001.1), *Hibiscus syriacus* (XP_039051919.1) and *Arabidopsis thaliana* (AT5G51190), and the AP2/ERF domain contains conserved YRG and RAYD elements, which may play a key role in DNA binding and protein interactions (Figure 1B). Using the CDD website (Marchler-Bauer et al., 2009) to predict the conserved domain of *GauERF105*, the results show that the amino acid sequence contains a typical AP2 domain at positions 58-125aa (Figure1C). Phylogenetic analysis confirms that *GauERF105* had a high similarity with different cotton species but it is more closely related to *Hibiscus syriacus* than *Herrania umbratical* and *Arabidopsis thaliana* (Figure 1D).

**Figure 1.**
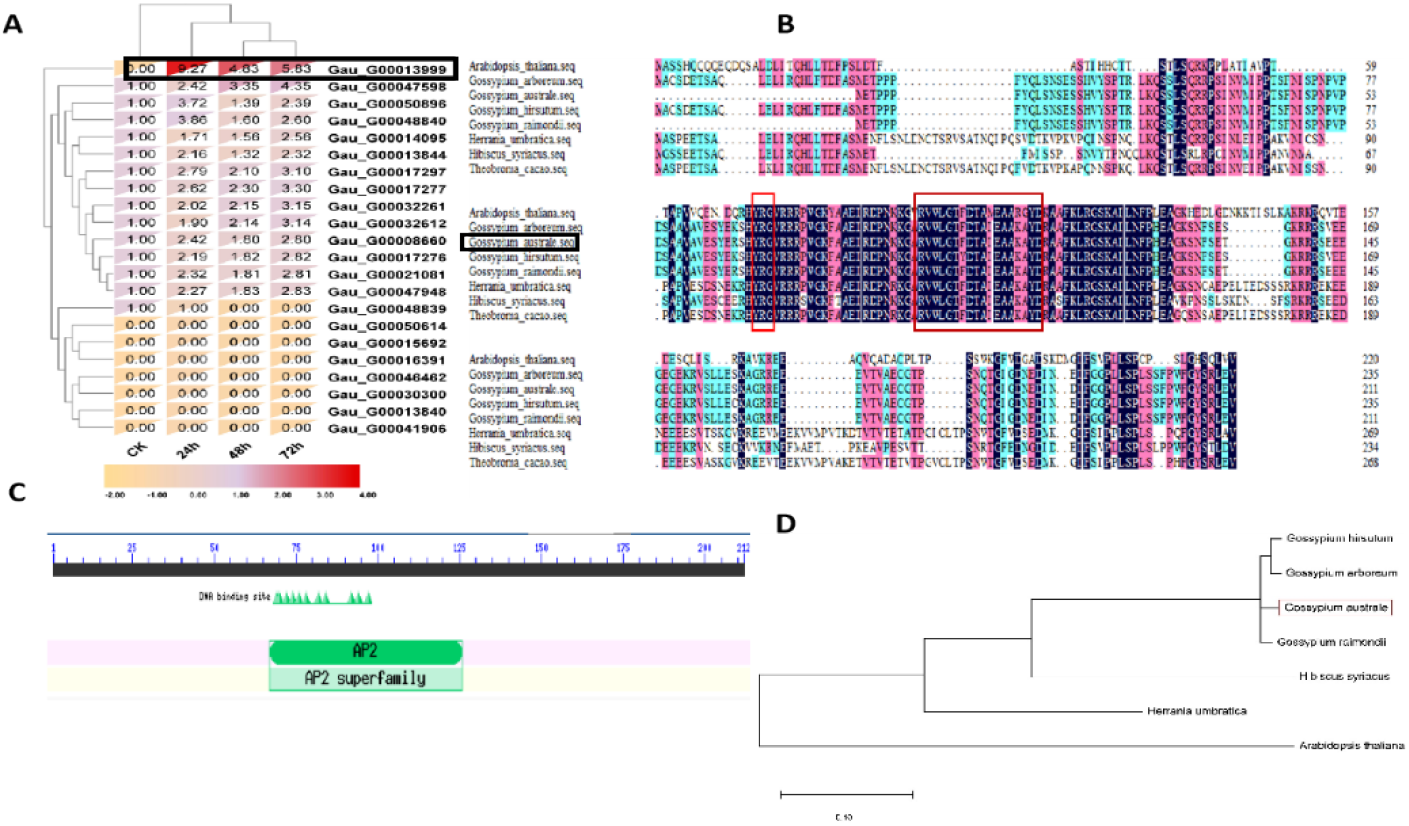
Cloning and Sequence Analysis of *GauERF105* **A-** Gene structure analysis of *GauERF105* (Gau_G00013999), **B-** Multiple sequence alignment of *GauERF105*, **C-** Prediction of conserved domain for *GauERF105*, **D-** Phylogenetic analysis of *GauERF105*

### 3.2 Expression analysis of *GauERF105* under *Verticillium dahliae* stress

Expression patterns of *GauERF105* have been evaluated in the leaves of *Gossypium australe* at 0h, 3h, 6h, 12h and, 24h after applying *Verticillium dahliae* in order to confirm the role of selected gene in response to *Verticillium dahliae*. It was observed that after inoculation, the expression level of *GauERF105* gradually increased from 0h to 24h post inoculation (Figure 4). Highest gene expression was observed at 24h post inoculation. The results showed that *GauERF105* plays a critical role in response to *Verticillium dahlia* attack and it might be the key candidate involved in Verticillium wilt resistance.

### 3.3 Overexpression of *GauERF105* enhances the resistance of Arabidopsis in response to Verticillium wilt attack

To validate the role of *GauERF105* in Verticillium wilt resistance we constructed an overexpression vector “PBI121-*GauERF105*” and transform into *Arabidopsis thaliana. Arabidopsis thaliana* was infiltrated using floral method. We selected positive seedlings and screened them on the MS solid medium containing kanamycin until the T_3_ generation where homozygous lines were screened. Samples from eleven positive T_3_ generation plants were collected to analyze the expression levels of *GauERF105*. Based on the results of RT-qPCR and agarose gel electrophoresis, we selected OE1 and OE2 for subsequent experiments **(Figure 3A, B)**.

**Figure 2:**
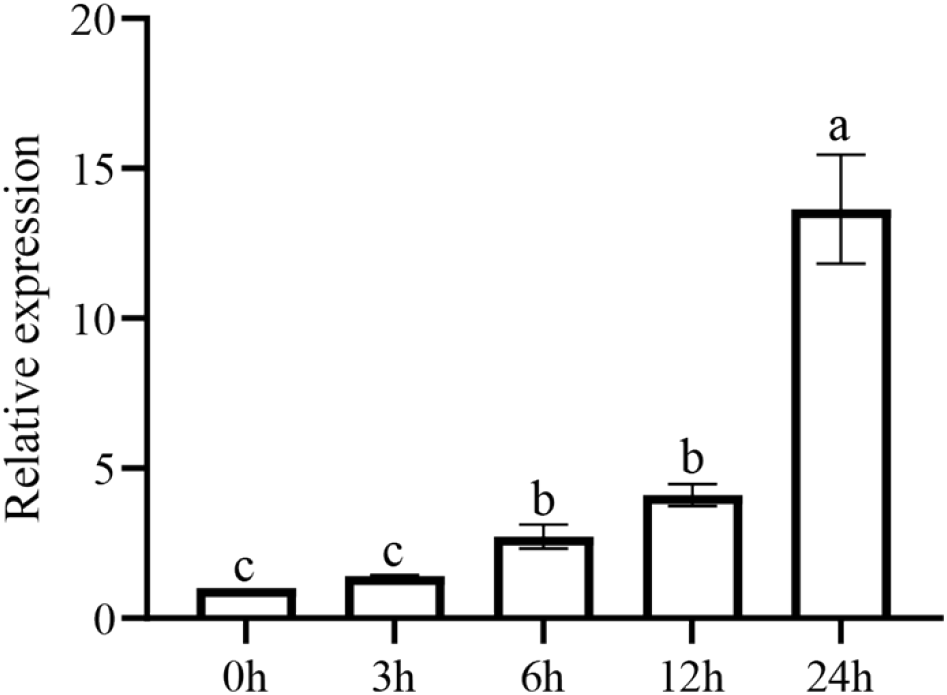
Expression pattern of *GauERF105* via RT-qPCR at 0, 3, 6, 12 and 24 hours post *Verticillium dahlia* inoculation.

**Figure 3:**
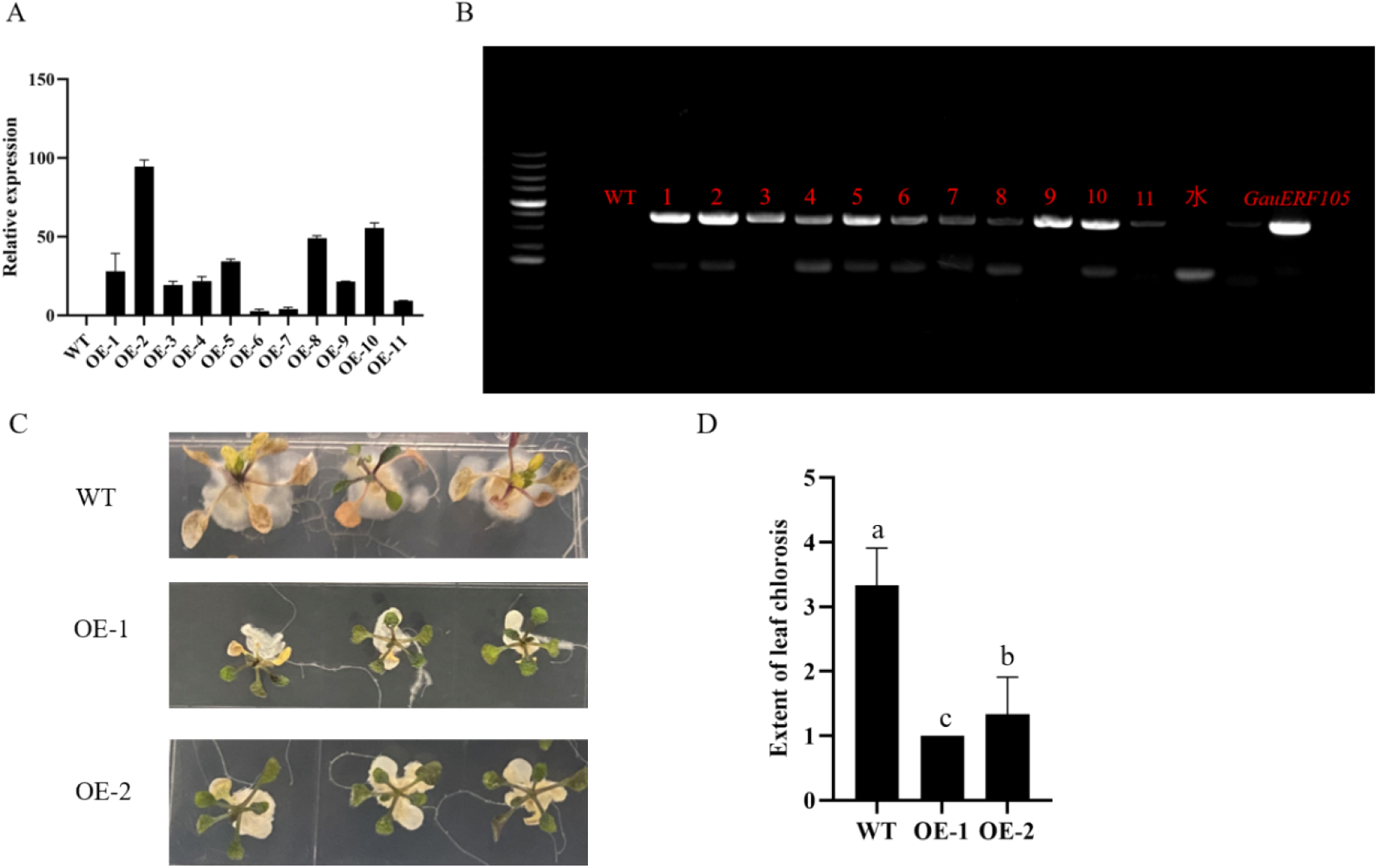
Overexpression of GauERF105 enhances the resistance of Arabidopsis in response to Verticillium wilt attack **A-** Relative expression of *GauERF105* in the overexpressed lines, **B-** Polymerase chain reaction (PCR) to confirm 639 bp coding sequence (CDS) integration in transformed T2 generation, number 1–11 transgenic lines, **C-** Transgenic lines and wild type under normal and diseased conditions, **D-** Extent of leaf chlorosis in wildtype and overexpressed lines after fungal inoculation.

The seeds of OE1 and OE2 lines were spot planted on MS medium, a small amount of *Verticillium dahliae* was added to MS medium, and the growth of plants was observed **(Figure 3C)**. Results indicated that WT showed a more sensitive phenotype to *Verticillium dahliae* as compared to the transgenic lines “OE1 and OE2”. In addition, the degree of WT plants resistance towards verticillium wilt was higher than that of transgenic lines, which indicates that the overexpression of *GauERF105* gene enhances the plant’s resistance to Verticillium wilt. We also planted wild-type Arabidopsis and transgenic lines in nutrient soil and observed the symptoms after inoculation. After overexpressing the *GauERF105* gene in Arabidopsis, the plant ability to resist Verticillium wilt was significantly enhanced, which is consistent with the phenotype of Arabidopsis grown on MS plates. The disease index was further counted by using the formula already mentioned in the methodology section, and a quantitative experiment of *Verticillium dahliae* was carried out, we observed results were consistent with that of phenotypic observations (**Figure 3D)**.

### 3.4 DAB and trypan blue staining

The accumulation of ROS represents the oxidative damage which occurs due to the stress caused by *Verticillium dahliae* attack. After 72 hours of inoculation leaves were taken for DAB staining. Wild-type Arabidopsis and transgenic lines both started to accumulate ROS but the leaves of WT plants are affected more as compared to transgenic lines, indicating that the overexpression of *GauERF105* reduced the damage caused by *Verticillium dahliae* attack.

Due to the damage of *Verticillium dahliae*, a large number of dead cells were produced in the plants. Therefore, the wild-type plants are stained darker. At the same time, the staining area of the wild-type *Arabidopsis thaliana* after inoculation was significantly larger than that of the transgenic *Arabidopsis thaliana*, which indicated that the damage of *Verticillium dahliae* to plants was greatly reduced after the *GauERF105* gene was overexpressed **Figure 4**.

**Figure 4:**
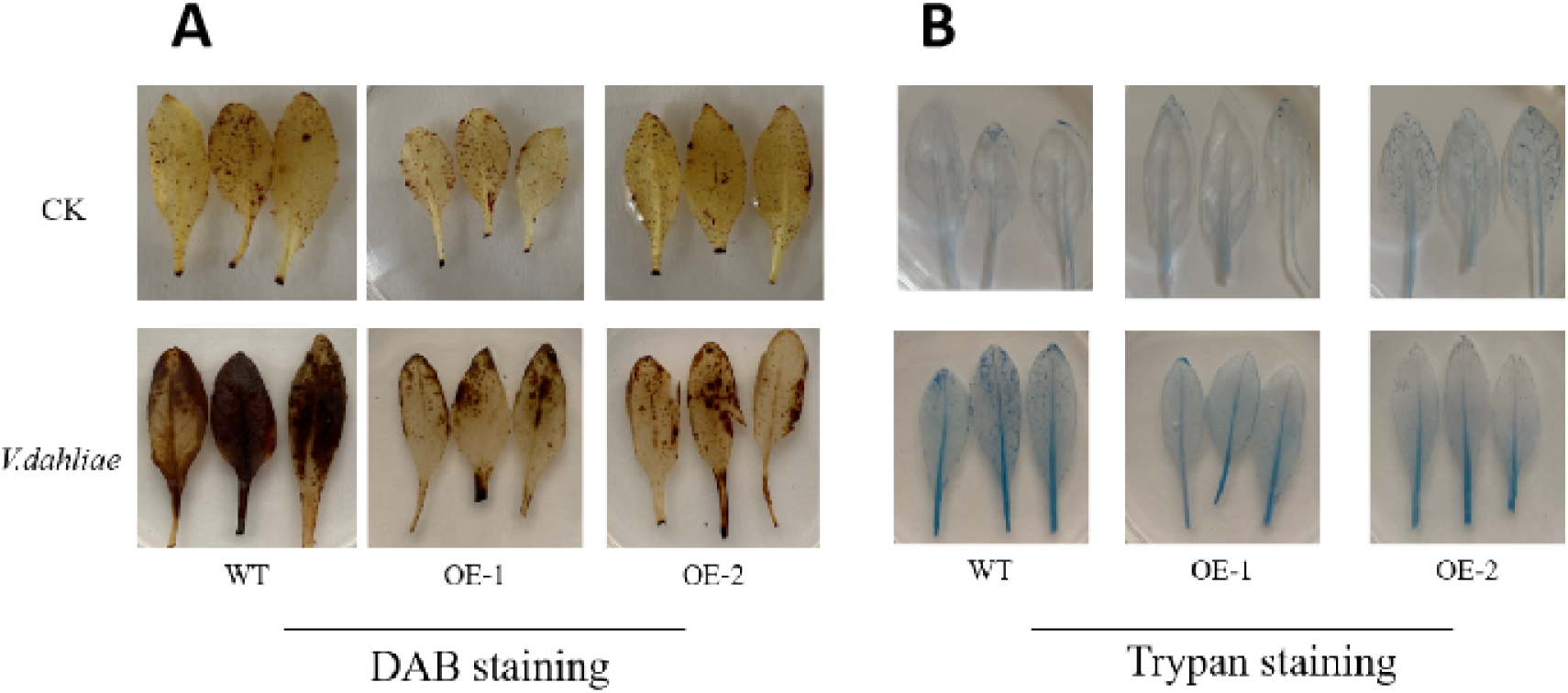
DAB and trypan blue staining **A**-DAB staining to estimate the damage on Arabidopsis leaves after fungal inoculation, **B-** DAB staining to estimate the damage on Arabidopsis leaves after fungal inoculation

### 3.5 Silencing of *GauERF105* gene decreases the resistance against Verticillium wilt in cotton

In order to verify the function of *GauERF105* in response to Verticillium wilt attack in cotton, virus-induced gene silencing was used to silence the homologous gene *GhERF105* in upland cotton **(Figure 5)**. About 13 days after VIGS, the cotton leaves were injected with TRV:PDS bacteria, chlorosis started and an albino phenotype was observed, which proved that the VIGS system was established successfully and the results were accurate for further experiment **(Figure 5A)**. The qRT-PCR results showed that the expression level of *GhERF105* gene was significantly lower in silenced plants as compared to WT and TRV:00, indicating that *GhERF105* gene is accurately silent.

**Figure 5:**
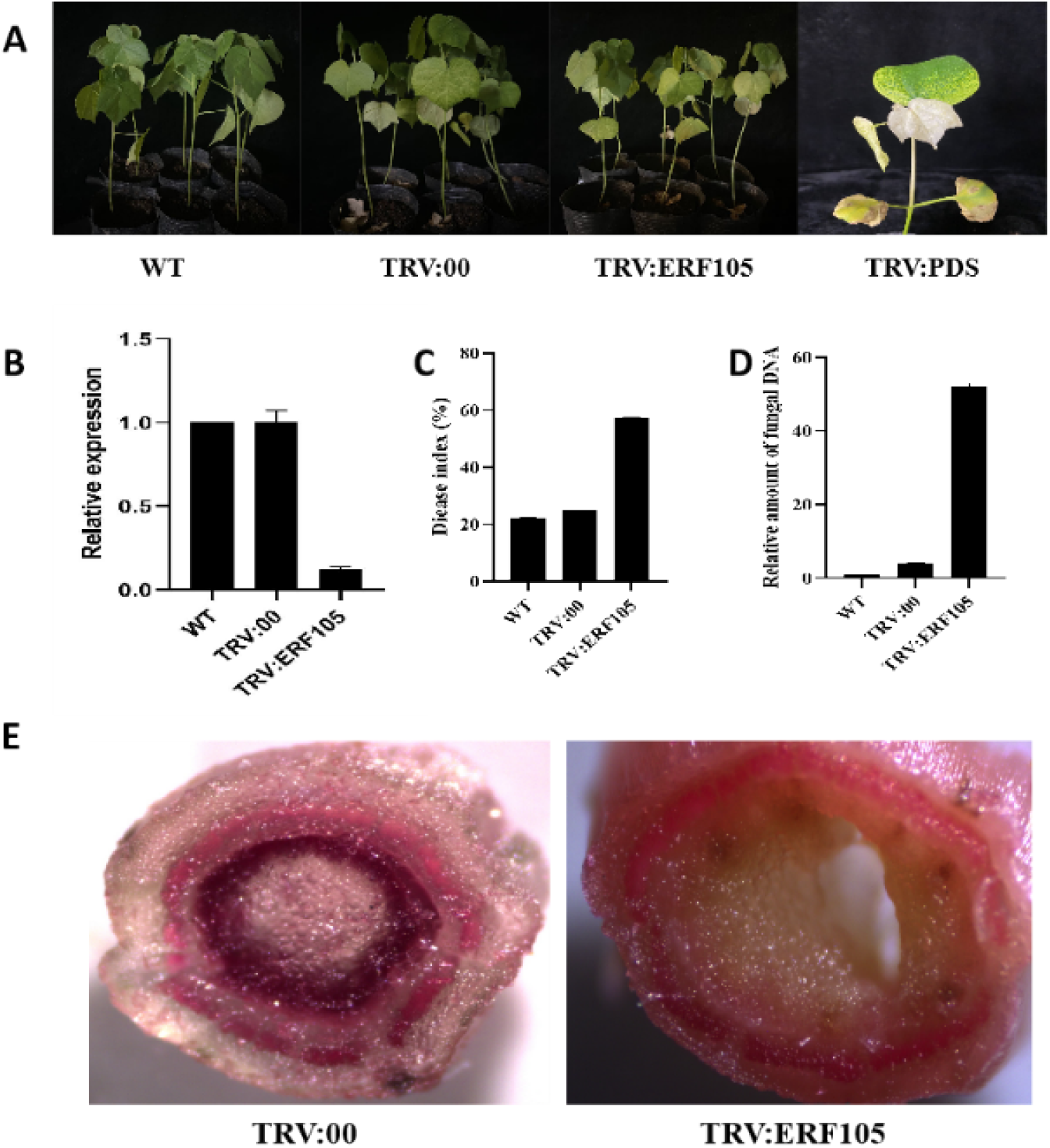
Silencing of *GauERF105* gene decreases the resistance against Verticillium wilt in cotton **A-** Representative images of WT, Positive Control, and VIGS plants, **B-** Relative expression of *GhERF105* in WT, TRV:00 and TRV:GhERF105, **C-** Disease index (%) WT, TRV:00 and TRV: GhERF105, **D-** Relative amount of fungal DNA in WT, TRV:00 and TRV:GhERF105, **E-** Histochemical staining of cotton stem lignin. Bars show standard error. WT: Wild type, TRV:00 Positive control, TRV:GhERF105, the VIGS plants.

Further, wildtype plants, empty vector plants and silent plants were inoculated with *Verticillium dahliae*, and the phenotype was observed after 25 days of inoculation **(Figure 5B)**. Compared with the control plants, the leaves of the silent plants turned yellow, wilted, and even fell off, and the disease index of the silent plants was also significantly higher. The degree of infection in silent plants was severe as compared to control plants **(Figure 5C)**. In addition, the leaves of WT, TRV:00 and TRV:GhERF105 plants were quantified for *Verticillium dahliae*. The expression level of *Verticillium dahliae* in the silenced target gene plants were significantly higher than that of the control plants, which was consistent with the results of the previous disease and disease index investigation **(Figure 5D)**. We sterilized the cotton stems after inoculation and cultured them in a PDA solid medium. The number of *Verticillium dahliae* in the TRV:00 plants were significantly smaller than that of the TRV:ERF105 plant, indicating that *Verticillium dahliae*. This indicates that silencing of *GhERF105* gene weakens the plant’s resistance to Verticillium wilt attack and make the plant more vulnerable to damage. Cotton lignin dying results showed that silent plants have inhibition of lignin as compared to wildtype and non-silent plants **(Figure 5E)**. Thus, proving the role of *GhERF105* in VW resistance in cotton.

### 3.6 Expression of disease-resistant marker genes in Cotton

In order to further analyze the regulatory role of *GauERF105* gene in the process of plant disease resistance, we further screened the expression of some disease-resistant pathway related genes in Arabidopsis and cotton (**Figure 6 A, B, C)**. The results showed that when the plants were inoculated with *Verticillium dahliae*, the expression of PRs increased; the expression of *AtPDF1*.*2, AtPR3* and *AtPR4* genes in the two *GauERF105* gene-transformed transgene lines OE-1 and OE-2 lines was significantly higher than that in the wildtype. In VIGS plants, the expression of PRs genes in the silenced plants was significantly downregulated. It further illustrates that the *GauERF105* gene can activate hormone-related pathways to participate in plant disease resistance (**Figure 6 D, E, F)**.

**Figure 6:**
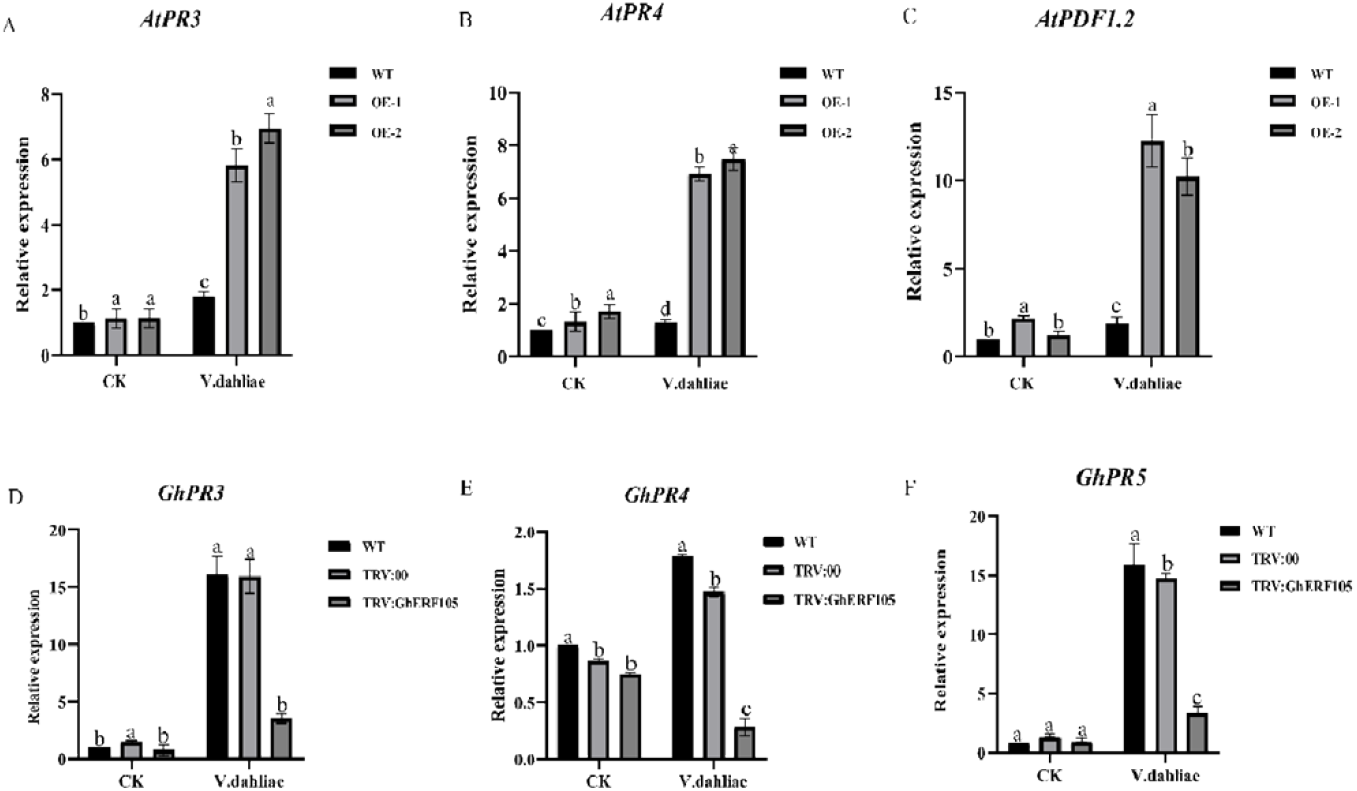
Expression of disease resistant marker genes in Transgenic Arabidopsis and Cotton.

## 4 Discussion

Plants opt a series of defense mechanisms after being invaded by pathogens. As one of the largest transcription factor families in plants, ERFs participate in the regulation of plant disease resistance.

ERF transcription factors positively activate the expression of genes related to plant resistance to pathogens or regulate the accumulation of some secondary metabolites in plants, thereby enhancing resistance to pests and diseases (SHAO, SHI, ZHANG, & LANG, 2021). In the interaction network of plant immune response, SA and JA/ET have cross-effects (Guo et al., 2016). Studies have shown that the SA pathway is involved in regulating the defense of plants against living vegetative pathogens, and JA and ET signal transduction are considered to be effective against pathogen attack. Necrotrophic pathogens such as *B. cinerea* and *F. oxysporum* are more effective (Derksen, Rampitsch, & Daayf, 2013). In this study, the *GauERF105* gene was screened and cloned from *G. australe*. Here we overexpressed *GauERF105* in Arabidopsis and checked the expression levels of AtPR3, AtPR4 and AtPDF1.2 genes in the transgenic lines OE-1 and OE-2. We observed that, the expressions were significantly higher in transgene lines as compared to wild-type Arabidopsis **(Figure 3)**, which shows that when the expression level of *GauERF105* gene increases, the expression level of its downstream genes also increases; DAB staining results show that the staining degree of *GauERF105* in transgenic Arabidopsis leaves is lighter than that of wild-type Arabidopsis. These result shows that overexpression of the *GauERF105* gene reduces the oxidative damage of pathogens to plants and improves the resistance of Arabidopsis against Verticillium wilt. PR1 and PR5 are the downstream genes of the SA pathway, and PR3 and PR4 are the downstream genes of the ET/JA pathway. The promoter regions of these disease related proteins have GCC-box. ERF transcription factors activate the expression of downstream defense genes by binding to GCC-box. So, as to enhance the plant’s resistance to diseases and insects attacks. After overexpression of the potato *StERF94* transcription factor, the expression of PRs-related genes increased, thereby enhancing the potato’s resistance to *Fusarium oxysporum* (Charfeddine, Samet, Charfeddine, Bouaziz, & Bouzid, 2019). Wang et al. (Wang, Liu, & Wang, 2020) overexpressed the *VqERF112, VqERF114* and *VqERF072* genes in Arabidopsis, and activated the SA signal-related genes AtNPR1 and AtPR1 and JA/ET signal-related genes AtPDF1.2, AtLOX3, AtPR3 and AtPR4, thereby enhancing the expression of Arabidopsis resistance to *Pst-DC3000* and *B. cinerea*.

We used VIGS to verify the role of *GauERF105* homologous gene *GhERF105* in upland cotton in the disease-resistant variety Zhongzhimian No. 2, after inoculation with *Verticillium dahliae*. The silent plants turned yellow, wilted, or even died, compared with the control plants. After silencing *GauERF105* gene, plants are more sensitive to Verticillium wilt **(Figure 5)**. Cotton disease index survey **(Figure 5C)**, lignin staining **(Figure 5E)**, *Verticillium dahliae* recovery culture, and quantitative experiment of *Verticillium dahliae* **(Figure 5D)** showed that the silencing of *GauERF105* gene, weaken the defense ability of plants against pathogens. Compared with the control, the expressions of GhPR3, GhPR4 and GhPR5 were significantly downregulated in the silent plants **(Figure 6 D, E, F)**, which indicated that when the expression of *GhERF105* gene was interfered, the expression of its downstream genes was inhibited, which weakened the plant’s disease resistance. Compared with non-inoculated plants, the expression levels of disease-related protein genes in cotton or *Arabidopsis thaliana* were increased, indicating that the plants activated SA, ET/JA, and other hormone transmission pathways after being subjected to biological stress., In order to participate in the defense response of plants.

The above results indicate that *GauERF105* acts as a positive regulator of plant Verticillium wilt resistance in both the model plant Arabidopsis and cotton.

## 4 Conclusions

Verticillium wilt attack on cotton are very severe in China and results in more and more yield loss every year. Therefore, it is an utmost requirement to have disease resistant cotton varieties. For that purpose, we need the candidate genes responsible for disease resistant especially VW in cotton. Here, we selected and screened that *GauERF105* gene from *Gossypium Australe* in order to verify its potential role against verticillium wilt attack in cotton and Arabidopsis. We performed overexpression experiments in Arabidopsis and VIGS in cotton. Our results indicated that overexpression of *GauERF105* increases the disease resistance ability in Arabidopsis and by silencing *GauERF105*, results in a decrease in defense and resistance. RT-qPCR, Trypan blue, DAB and lignin staining also validates our findings and hence it is proved that *GauERF105* is a truer candidate gene for resistance against VW attack in cotton. This gene can be used for further breeding programs to create the disease resistance against VW attack.

## Authors’ statement

W.Y.Q, M.J.U, and Y.X conducted the experiment and wrote the manuscript. R. O. M, T.G.M and M. L.S, Z.J assisted in data analysis. K.W., X.C, YH, YW, Z.Z., and F.L. revised the manuscript. All authors reviewed and approved the final manuscript.

## Acknowledgments

We are very much thankful to the Institute of Cotton Research, Chinese Academy of Agricultural Sciences and to our laboratory for providing the full supply and support during the experiment.

